# Quantification of Mouse Total Body Surface Area: Implications for Preclinical Burn Research

**DOI:** 10.64898/2026.04.30.722020

**Authors:** Abby Barlow, Martin Morales, Mary Barre, Meagan Kingren, Craig Porter

## Abstract

Clinically, burn severity is reported as the size (and depth) of burn wounds relative to total body surface area (TBSA). This nomenclature is also often used in rodent models of burns. Accordingly, accurate determination and reporting of rodent TBSA is required to ensure the rigor and reproducibility of preclinical burn research. Rodent TBSA is typically estimated indirectly as a function of body mass. Further, empirical quantification of rodent TBSA through pelt dissection does not consider differences in rodent and human anatomy, making comparison of relative burn size in rodents and humans a challenge. Here, we compared commonly used approaches to directly determine or indirectly estimate rodent TBSA to demonstrate the impact different approaches can have on the calculation of relative burn size.

A total of n=48 C57BL/6J background mice (55% male) ranging from 4 to 45 weeks of age and 17 to 40 grams were used. Mice were weighed prior to euthanasia. After euthanasia, mouse length was measured from the nose to anus. Mice were then placed into clear polypropylene sheet protectors (21.6 x 27.9 cm) to trace the areas of both the dorsal and ventral surfaces as well as all four limbs (dorsal-ventral (DV) tracing). Next, the pelt was carefully excised from the body through cutting a lateral line from the mouth to the genitalia, then again proximally to distally on all four limbs. The pelt was gently placed on a sheet protector and traced when both relaxed and stretched. The ears and tail were removed and traced separately. Photographs were taken of all tracings next to a ruler for scale and analyzed in ImageJ.

Stretched pelt measurements of TBSA were 34% (79.4±7.6 vs. 57.5±7.5 cm^2^, P<0.001) and 30% (70.6±10.9 vs. 52.7±8.1 cm^2^, P<0.001) greater than relaxed pelt TBSA measurements in male and female mice respectively. TBSA estimated by DV tracing was 9% greater in males (62.5±10.9 vs. 57.5±7.5 cm^2^) and 15% in females (60.6±12.3 vs. 52.7±8.1 cm^2^) compared to TBSA measurements made on relaxed pelts. Accordingly, empirically derived Meeh constants (*k*) from DV tracing were greater than those derived from relaxed pelt measurements for both males (7.14±0.59 vs. 6.58±0.72) and females (7.72±0.58 vs. 6.78±0.80). In contrast *k* values derived from stretched pelt measures of TBSA were significantly greater than those determined in relaxed pelts for males (8.91±0.87 vs. 6.58±0.72, P<0.001) and females (8.85±1.25 vs. 6.78±0.80, P>0.001). The combined ears and tail represent approximately 7% and 8% of the TBSA measured by the relaxed pelt approach, respectively. Exclusion of the tail and ears from the calculated TBSA results in derived *k* values that are ∼16-17% lower.

The approach used to determine TBSA in mice significantly influences measured areas and thus derived *k* values. We suggest that stretching the pelt prior to tracing inflates TBSA values, where measurements made from relaxed pelts or by DV tracing likely provide more accurate estimates of actual TBSA. Further, exclusion of the tail and ears (the latter of which is not typically considered in estimates of TBSA in humans) may be a useful approach relating relative burn sizes of mice to those of humans.

## INTRODUCTION

Burns represent a leading cause of non-fatal trauma that can result in a prolonged metabolic stress response (1). This stress response is associated with long-term morbidity and is typically a function of burn severity. Burn injury severity relates to the size (and depth) of burn wounds relative to the patient’s total body surface area (TBSA), where the relative size of full thickness burn wounds correlate with the degree of burn-induced hypermetabolism (2) and cachexia (3, 4). Relative burn size is also frequently used as a primary indicator of injury severity in rodent models of burn injury. Rodent TBSA is typically estimated indirectly as a function of body mass (BM) by way of the Meeh formula (5), where TBSA is equal to *k*.BM^2/3^. While *k* is an empirically determined species specific coefficient, that is also influenced by strain and body composition in mice (6), historical values for *k* are often used in place of those determined directly given that direct determination of TBSA and thus calculation of *k* can be tedious.

Empirical determination of TBSA (and thus *k*) requires careful dissection of the pelt, which is time consuming and can be technically challenging. Indeed, published *k* values determined in rats range from ∼7 to 12.56 (7-12), which can result in estimated TBSA’s that vary by as much as 32%, which in the context of burn injury, would significantly under or overestimate relative burn size. Further, empirical quantification of rodent TBSA through pelt dissection often does not account for the wide-spread presence of the *panniculus carnosis* muscle, which encompasses most of the rodent torso (13), or for critical differences in rodent and human anatomy, making comparison of estimated relative burn size in rodents and humans challenging.

The objectives of this work were to (i) compare commonly used approaches for direct and indirect determination of rodent TBSA and (ii) to determine the contributions of key anatomical structures to mouse TBSA in order to devise a more translational approach to TBSA estimation in rodents. Our data suggest that stretching the pelt prior to tracing inflates TBSA values, where measurements made from relaxed pelts or by DV tracing provide more accurate estimates of actual TBSA. Further, exclusion of the tail and ears (the latter of which is not typically considered in estimates of TBSA in humans) may be a useful approach when relating relative burn sizes of mice to those of humans.

## METHODS

### Animals

A total of n=49 C57BL/6J background mice (45% female) ranging from 4 to 45 weeks of age and 17 to 40 grams were used. Mice were housed at 24°C under a 12/12 light/dark cycle. All mice were surplus animals generated through colony breeding at our facility and were humanely euthanized in a rising concentration of CO_2_ prior to use in the current study.

### Determination of Total Body Surface Area

#### Dorsal-Ventral Tracing

After euthanasia, the length of each mouse was measured from the nose to anus and recorded. Mice were then placed into a clear polypropylene sheet protector (21.6 x 27.9 cm) dorsal side up to trace the area of the dorsal surface. Then, mice were removed from the sheet protector and placed on top of the dorsal tracing, and the ventral surface was then traced (**Figures 1A to 1C**). The tail and each ear were individually traced upon dissection and the recorded areas double to account for both sides of the tail and ears. The area of the legs were traced while the mouse was in prone position (**Figure 1A-C**) and doubled to account for both sides of each limb.

**Figure 1.**
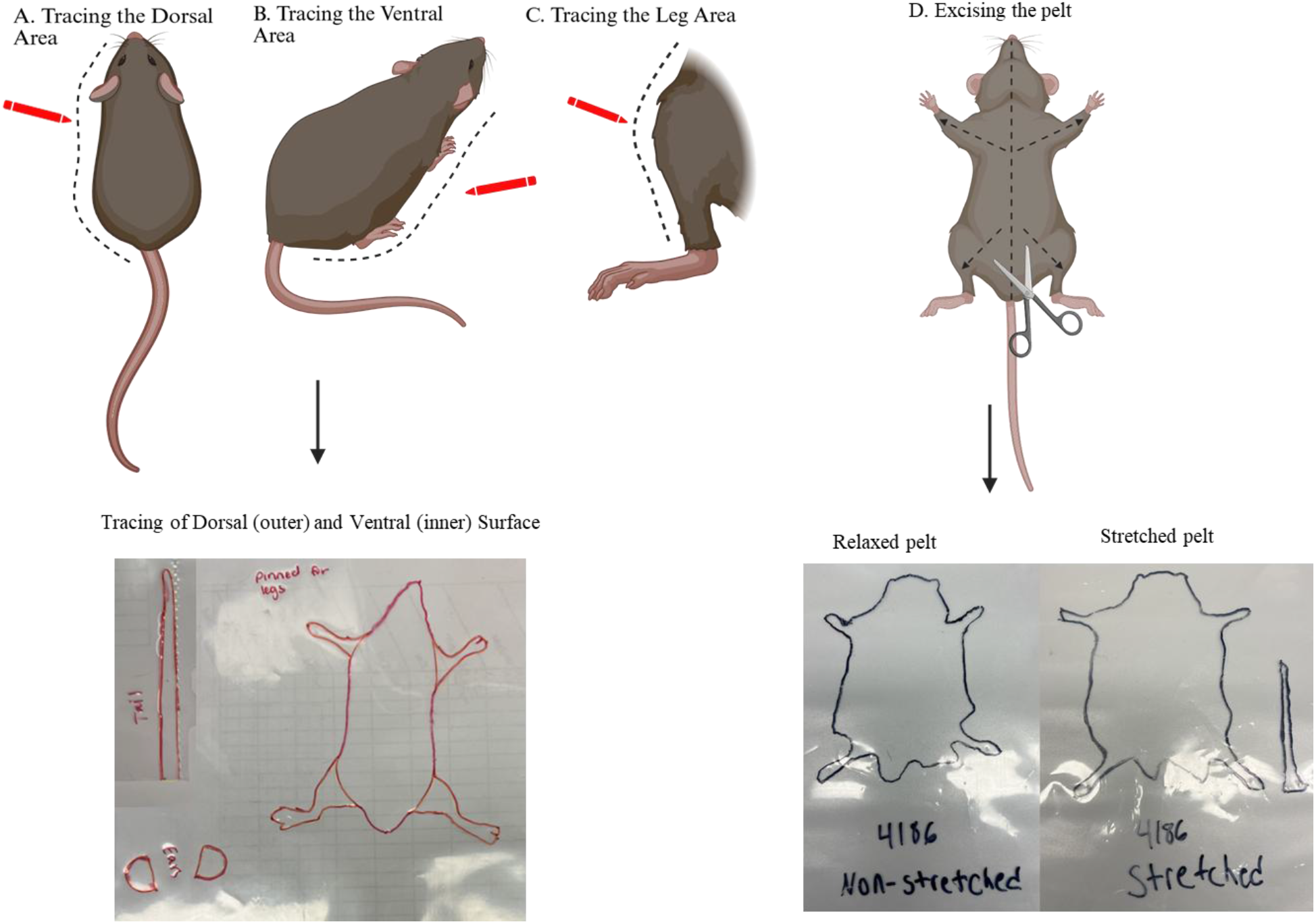
Methodological approach. Overview of approach for dorsal-ventral tracing approach (Panels A and B) where the legs (Panel C) as well as the tail are traced separately. Panel D depicts the dissection if the pelt away from the underlying tissue for tracing while placed on an acetate sheet while relaxed (non-stretched) and stretched.

#### Direct Measurement of the Pelt Area

To measure the area of the pelt, the pelt was carefully excised from the body through cutting a lateral line from the mouth to the genitalia, then again proximally to distally on all four legs (**Figure 1D**). The pelt was then excised from the body carefully starting with the trunk, then legs, then moving on to the head. To measure the relaxed pelt surface area, the pelt was gently placed on a sheet protector without stretching and traced around. For measuring the stretched surface area, the pelt was gently stretched and pinned on top of a sheet protector before tracing (**Figure 1D**). The ears and tail were excised and measured separately as described above.

#### Image Analysis

Photographs were taken of the tracings then uploaded into ImageJ. A ruler was placed in all images as scale reference, and the tracings subsequently measured using the Freehand Selection tool.

## RESULTS

### Animals

Demographic data for male and female mice are reported in **Table 1**. Data for the measured area of both ears, the tail, legs (all four limbs combined), the dorsal and ventral area (determined by tracing) and the relaxed and stretched pelt for male and female mice are reported in **Table 1**. Pelt values reported in **Table 1** do not include the tail or ears.

**Table 1.**
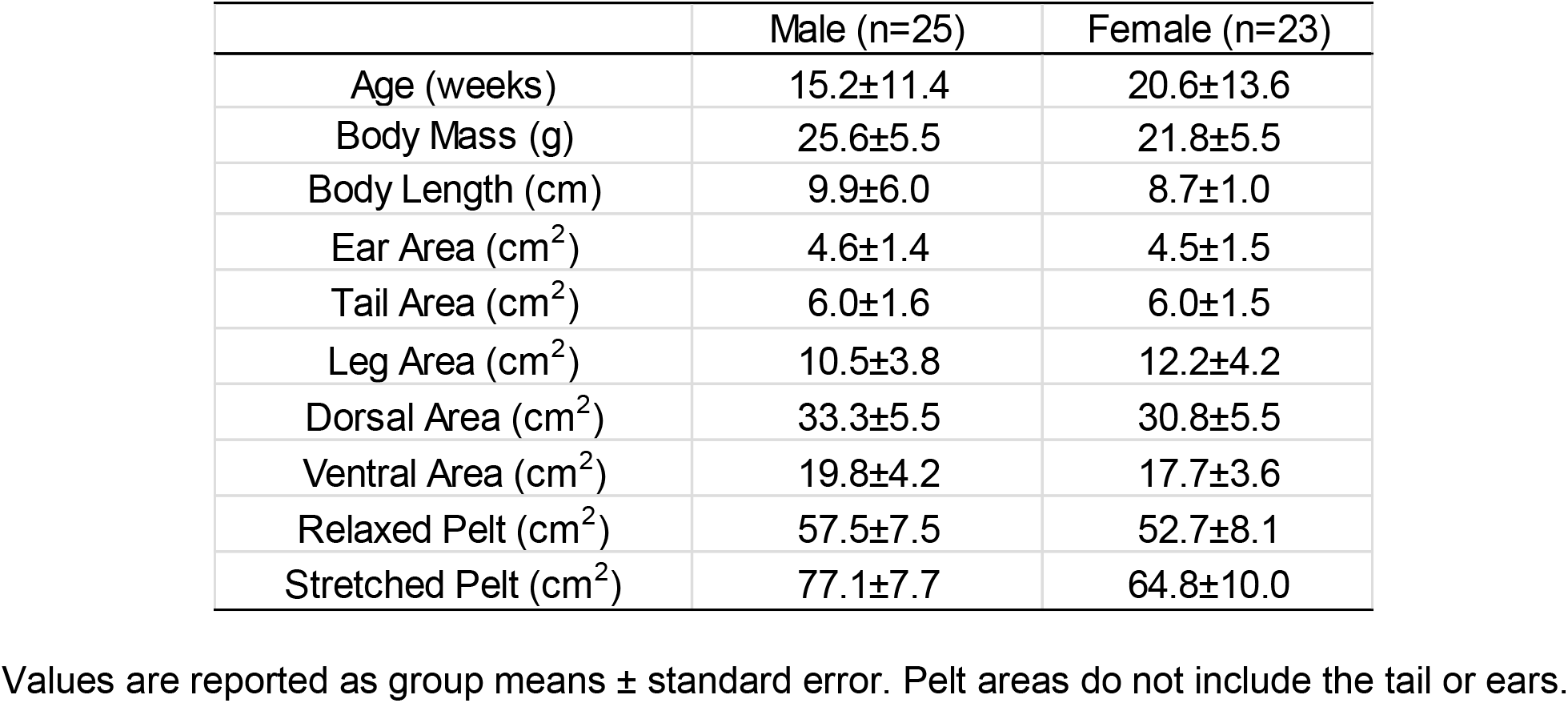
Animal Demographics.

### Determination of TBSA

TBSA data derived for DV-tracing or measurement of relaxed and stretched pelts is reported in **Table 2** along with corresponding Meeh constants (*k*). Calculating TBSA by summing the dorsal and ventral areas with the combined areas of the limbs was greater than the determination of TBSA by measurement of the relaxed pelt in both male (62.5±10.9 vs. 57.0±7.4 cm^2^, P=0.05) and female (60.6±10.5 vs. 52.7±8.1 cm^2^, P=0.02) mice. Accordingly, empirically derived *k* values from DV-tracing and measurement of the relaxed pelt were also greater in both males (7.14±0.59 vs. 6.58±0.72) and females (7.72±0.58 vs. 6.58±0.72).

**Table 2.**

|  | ♂ |  | ♀ |  |
| --- | --- | --- | --- | --- |
|  | BSA (cm<sup>2</sup>) | Meeh Constant (k) | BSA (cm<sup>2</sup>) | Meeh Constant (k) |
| DV Tracing | 62.5±10.9 | 7.14±0.59 | 60.6±12.3 | 7.72±0.58 |
| Pelt (relaxed) | 57.0±7.4 | 6.58±0.72 | 52.7±8.1 | 6.78±0.80 |
| Pelt (stretched) | 77.1±7.7 | 8.91±0.88 | 68.4±10.0 | 8.85±1.25 |

Values are reported as group means ± standard error. DV-sketch or pelt areas do not include the tail or ears.

Measured TBSA was 34% (79.4±7.6 vs. 57.5±7.5 cm^2^, P<0.001) and 30% (70.6±10.9 vs. 52.7±8.1 cm^2^, P<0.001) greater when determined in pelts that were stretched rather than relaxed for male and female mice, respectively (**Table 2**). This meant *k* values derived from stretched pelt measurement of TBSA were also ∼34% and 30% greater than those derived from the relaxed pelt measurement in males and females, respectively (**Table 2**).

### Correlation Between Body Mass and Body Surface Area

Total body surface area is often determined indirectly using body mass and the Meeh equation. While excision of the pelt is considered the gold standard approach for determination of TBSA, this procedure can be technically challenging, as is standardizing the pining of the pelt to measure its maximum area in the ‘stretched’ state. In our hands, the correlation between body mass and TBSA was highest when TBSA was determined by the dorsal-ventral tracing (r=0.87, P<0.001) versus the relaxed pelt (r=0.71, P<0.001) and stretched pelt (r=0.66, P<0.01) (**Figure 2**). Similarly, variance was lower when TBSA was determined by the dorsal-ventral tracing (*R*^*2*^=0.76) compared to when the relaxed pelt (*R*^*2*^=0.51) or stretched pelt (*R*^*2*^=0.43) was used to estimate TBSA (**Figure 2**).

**Figure 2.**
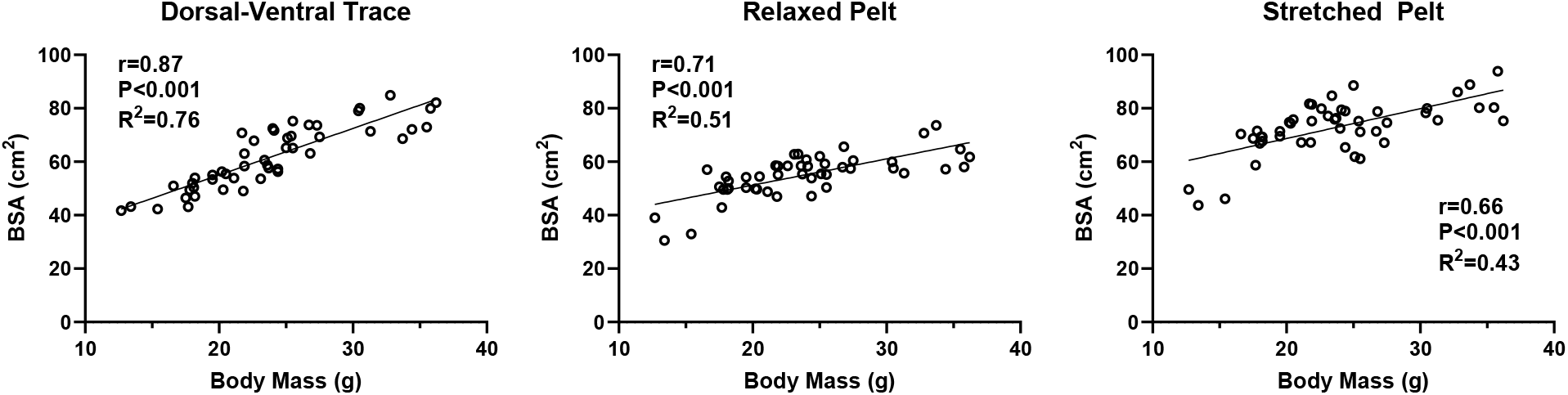
Impact of Total Body Surface Area Estimation of Relative Burn Size. Relationship between body mass and body surface area (BSA) determined by dorsal-ventral tracing, measuring the relaxed pelt, and by measuring the stretched pelt. Pearson correlation coefficients (*r*) and associated p values and the coefficient of determination (*R*^*2*^) for the linear regression curve are reported on each trace.

### Relating Mouse TBSA and Its Components to Humans

In the context of burn injury, estimation of TBSA in mice is useful for determining relative burn size in preclinical models. Since humans lack tails and the ears are typically not included in the estimation of human TBSA (particularly when estimating relative burn size in humans), we sought to determine the contribution of the ear and tail areas to the TBSA and derived *k* values in mice. The combined ears and tail areas of both male and female mouse were approximately 2.3 cm^2^ and 6.0 cm^2^, respectively (**Table 1**). When relating this to the measured area of the relaxed pelt, the area of the ears and tails account for approximately 7% and 8% of the TBSA, respectively. **Figure 3** shows graphical depictions of the relative contributions of key body parts to TBSA in mice and humans. Irrespective of whether the ears and tail are included, the trunk represents the vast majority of the TBSA of a mouse. This contrasts with humans, where the lower and upper limbs collectively represent over 50% of the TBSA of a human, as opposed to ∼16-17% in the mouse.

**Figure 3.**
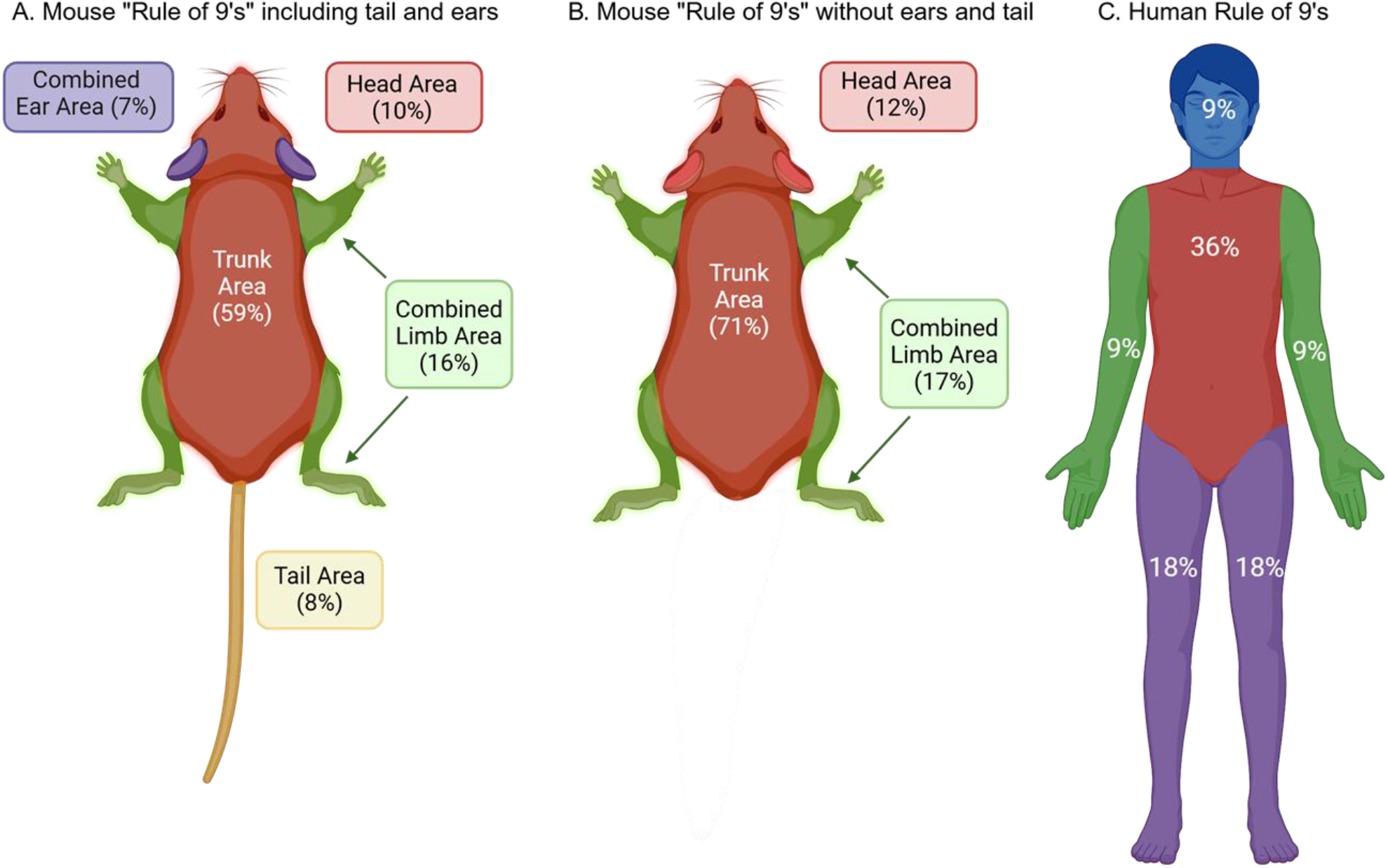
Determinants of Mouse and Human Body Surface Area. Overview of regional differences in the determinants of total body surface area (TBSA) in mice where the tail and ears are included (Panel A) or excluded (Panel B). Regional determinants of human TBSA by the rule of 9’s approach.

Anatomical differences influence the determinants of TBSA in mice and humans, therefore, when using mouse TBSA to estimate the relative size of a burn injury, inclusion of the area of the tail and ears markedly impacts relative burn size. For example, exclusion of the tail and ears results in a TBSA and derived *k* value that are ∼16% (P<0.001) lower for males and ∼17% lower for females (P<0.001) (**Table 3**). Thus, inclusion of the ears and tail when empirically determining mouse TBSA can significantly impact the estimation of relative burn size.

**Table 3.**

|  | ♂ |  | ♀ |  |
| --- | --- | --- | --- | --- |
|  | BSA (cm<sup>2</sup>) | Meeh Constant (k) | BSA (cm<sup>2</sup>) | Meeh Constant (k) |
| Pelt (+ ears and tail) | 67.7±7.9 | 7.80±0.78 | 63.1±8.7 | 8.15±1.00 |
| Pelt (- ears and tail) | 57.0±7.4 | 6.58±0.72 | 52.7±8.1 | 6.78±0.80 |

Values are reported as group means ± standard error. Areas reported are from relaxed pelt measurements.

### Impact of TBSA Determination of Estimation of Relative Burn Size

Figure 4 depicts a male (top) and female (bottom) mouse where TBSA has be calculated using four different Meeh constants derived from measuring TBSA by DV tracing, measuring the relaxed pelt, or measuring the stretched pelt with and without the inclusion of the ears and tail. The relative size of a burn wound (10 cm^2^ for males and 8 cm^2^ for females) is also calculated for each TBSA. This approach illustrates how the methodology used to determine TBSA and derive Meeh constants can markedly affect estimated TBSA and, in turn, the estimation of relative burn size, expressed as a percentage of the TBSA.

**Figure 4.**
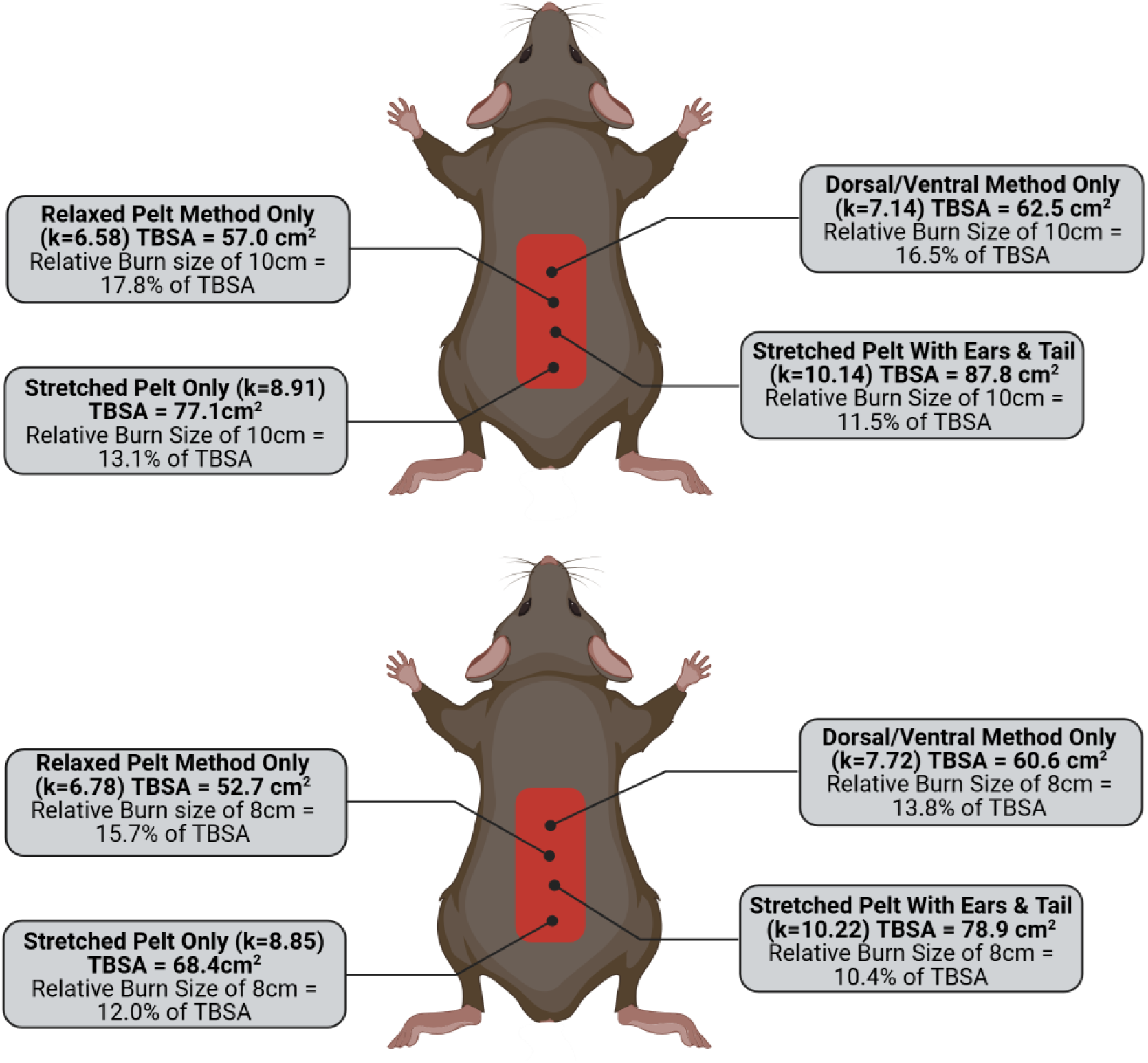
Impact of Total Body Surface Area Estimation of Relative Burn Size. Estimation of relative burn size (as a percentage of total body surface area (TBSA)) in male and female mice where TBSA was determined using dorsal-ventral tracing (including the limbs) the relaxed pelt, stretched pelt, and the stretched pelt (including the tail and ears).

For example, compared to using relaxed pelts (excluding the tail and ears) estimation of relative TBSA from a stretched pelt (including the tail and ears) results in relative burn sizes of 17.8±2.3% versus 11.5±1.1% in male mice (P<0.001) and 15.7±3.3% versus 10.4±1.8% in female mice (P<0.001). When comparing relative burn sizes calculated using TBSA values determined from relaxed pelts (excluding the tail and ears) were ∼36% (17.8±2.3% versus 13.1±1.4%, P<0.001) and ∼31% (15.7±3.3% versus 12.0±2.2%, P<0.001) than those calculated from TBSA values determined stretch pelts (excluding the tail and ears). While the DV tracing method resulted in greater TBSA values compared to those derived from measuring relaxed pelts, these two approaches resulted in relative burn sizes in male (17.8±2.3% versus 16.5±2.9%, P<0.08) and female (15.7±3.3% versus 13.8±2.9%, P<0.09). that were not significantly different.

## DISCUSSION

Clinically, the size of burn wounds relative to the patient’s TBSA is the primary means of reporting burn severity. This nomenclature is also used to indicate injury severity in rodent models of burn trauma. Rodent TBSA is typically determined indirectly via the Meeh equation, which relates TBSA to the two-thirds power of body mass (5), where TBSA = *k*.BM^2/3^. This approach necessitates the use of a previously published *k* value. Beyond differences in species, strain, and body composition (6) the methodological approach used to directly determine TBSA and subsequently calculate *k* may significantly impact these values. This means that TBSA determined indirectly via the Meeh equation may vary significantly across studies depending on the *k* value used in TBSA determination. Ultimately, this will result in the under or overestimation of relative burn size, making comparisons across studies challenging. Further, key differences in the relationship between rodent and human anatomy are not considered which comparing relative burn sizes, potentially limiting the translational impact of rodent models of burn trauma. In the current report, we determined the contribution of key anatomical structures to mouse TBSA and contrasted this to the human rule of 9’s approach (14) to devise a more translational and reproducible approach to TBSA estimation in rodents. Further, we compared commonly used approaches for direct and indirect determination of rodent TBSA, demonstrating that stretching and pining the pelt prior to tracing inflates TBSA values, where measurements made from relaxed pelts or by DV tracing provide more accurate estimates of actual TBSA.

We studied both male and female mice from a C57BL6/J background with a wide range in age (∼4-45 week) and body masses (17 to 40 grams). TBSA was estimated in all mice via DV tracing, prior to direct determination of TBSA via excision of the pelt. Corresponding Meeh constants (*k*) were then calculated for all three approaches. Estimation of TBSA by DV tracing has been reported to be straightforward and accurate approach in rats that does not require euthanasia (7). We found that summing the dorsal and ventral areas with the combined areas of the limbs (from DV tracing) resulted in an estimated TBSA that was significantly greater than direct determination of TBSA by measurement of the relaxed pelt (see **Table 2**). However, the DV sketching approach resulted in TBSA values that were more comparable to those derived from measuring the area of the relaxed pelt compared to those derived from measuring the area of the pinned stretched pelt.

Variations in reported *k* values for similar strains of rodent are likely due to the presence of the *panniculus carnosus* muscle. A thin layer of striated muscle, the *panniculus carnosus* attaches to both skin and fascia to allow twitching and contraction. While the *panniculus carnosus* may be present in discrete areas of most mammals including humans, loose skinned mammals such as mice have an extensive *panniculus carnosus* muscle layer encompassing the majority of the torso. The presence of the *panniculus carnosus* in excised pelts makes accurate determination of TBSA from pelt dimensions challenging since the pelt can potentially be stretched for measurement. Indeed, in the current report when pelts were stretched and pinned in place, the measured TBSA and calculated *k* values were 34% and 30% greater than relaxed male and female mice pelts, respectively. This significant inflation of TBSA and subsequently *k* values derived from these TBSA’s would lead to significant underestimation of relative burn size, since TBSA is the denominator and the fixed burn size is the numerator in this equation. Indeed, classic studies determining the TBSA of albino rats reported *k* values ranging from 12.53 to 7.47 (8-12). While many of these reports are ∼100 years old and provide varying degrees of methodological detail, upon closer examination it appears that the higher *k* value (11.38) were determined in pelts that were stretched and pinned prior to measurement. Studies that only report pinning (but not stretching) the pelt report lower *k* values (9.47). The lowest *k* value (7.53) for albino rats was reported after dipping rats in a collodion solution which was peeled off and measured once dried (9), thereby preventing any stretching or shrinking of the pelt once removed. Clearly, given the elasticity of the rodent pelt, approaches used to dissect off the pelt and pin it for measurement have the potential to have a significant impact on measured TBSA. Our current data in mice are in agreement with this, where *k* values calculated from TBSA measurements in relaxed and stretched pelts were markedly different in both male (6.58 versus 8.91) and female (6.95 versus 8.85) mice. Our current TBSA and *k* value data for stretched pelts are in line with those previously published for are range of mouse strains (6).

While excision of the pelt is considered the gold standard approach for determination of TBSA, this procedure can be technically challenging, and as illustrated above, where the amount of tension placed on the pelt when performing measurements is difficult to standardize and can significantly alter the measured TBSA (7). In our hands, the correlation between body mass and TBSA was highest when TBSA was determined by DV tracing (r=0.87) compared to either the relaxed pelt (r=0.71) and stretched pelt (r=0.66) approaches. Similarly, variance was lowest when TBSA was determined by DV tracing (*R*^*2*^=0.76) compared relaxed (*R*^*2*^=0.51) or stretched pelt (*R*^*2*^=0.43) approaches. This further suggests that both the dissection of the pelt and then its handling prior to measurement introduce error and variation. From a practical standpoint, this data indicates that the DV tracing approach is perhaps the most appropriate for researchers to use moving forward in terms of reducing variability across studies while ensuring measurement accuracy.

The determination of mouse TBSA can be useful in estimating relative burns sizes in preclinical models. However, it is important to consider some of the key differences in rodent and human anatomy and how they contribute to TBSA. When contrasting the determinants of mouse TBSA to human TBSA (using the rule of 9s (14)) it is clear that the torso represents the vast majority of the TBSA of a mouse. This is in contrast to humans, where the both the lower and upper limbs combined account for over 50% of the TBSA (as opposed to ∼16-17% in the mouse). Since humans lack tails and the ears are typically not included in the estimation of TBSA in humans, particularly when estimating relative burn size in humans, we also determined the relative contributions of both the ears and tail to TBSA and derived *k* values. In relation to the measured area of the relaxed pelt, the ears and tails account for approximately 7% and 8% of the TBSA, respectively (see **Figure 2**). Pertaining to the use of mouse TBSA in estimating the relative size of a burn wound, inclusion of the tail and ears significantly impacts relative burn size. For example, exclusion of the tail and ears results in a TBSA and derived *k* value that are ∼15% and ∼20% greater for male and females, respectively (see **Table 3**). Accordingly, inclusion of the ears and tail when estimating TBSA has the potential to significantly impact relative burn size estimation. Given that the ears are not included in the estimation of human TBSA via the rule of 9s – and that humans lack tails – we suggest that the omission of the tail and ears when estimating mouse TBSA represents a reasonable approach to making mouse TBSA more relatable to that of humans.

Our current data underscore two key points: (i) stretching the pelt prior to measurement introduces significant variation in measured TBSA and (ii) that inclusion of the ears and tail in mouse TBSA measurements inflate TBSA and influences relative wound size calculation. To illustrate these points further, we estimated relative wound size for a standardized burn wound ((10 cm^2^ for males and 8 cm^2^ for females) using TBSA values that were calculated from four different Meeh constants derived from dorsal-ventral tracing, measurement of the relaxed pelt, measurement of the stretched pelt, and finally from measurement of the stretched pelt including both the ears and tail (see **Figure 3**). TBSA measurements derived from stretched pelts versus TBSA derived from measurement of the relaxed pelt resulted in relative burn sizes that were 36% greater in males (17.8% versus 13.1%) and 21% greater in females (15.7% versus 12.0%). TBSA calculated from a stretched pelt alone versus a stretched pelt including the tail and ears results in relative burn sizes of 17.8% versus 11.5% in males and 15.7% vs ∼10.4% in females, respectively. The examples depicted in Figure 4 further underscore how the methodological approach used to determine TBSA can influence not only the measured TBSA and associated *k* value, but also the estimation of relative burn size.

Historically, there has been a lack of rigor in how rodent TBSA and relative wound size have been reported in the burn literature (15), owing in part to the use of varied *k* values to estimate animal TBSA. Our current data clearly demonstrate that the approach used to determine TBSA in mice significantly influences the measured body surface area and thus derived *k* values. Our data clearly demonstrates that stretching the pelt prior to tracing inflates TBSA values. We posit that measurements of TBSA made from relaxed pelts or by DV tracing provide more accurate estimates of actual TBSA. Given the relative ease associated with DV tracing and the tighter correlation seen between body mass and estimated TBSA, this approach may be the more straightforward to adopt broadly across the preclinical burn research field. Further, we suggest that exclusion of the tail and ears (the latter of which is not typically considered in estimates of TBSA in humans) may be a useful approach when relating relative burn sizes of mice to those of humans.

## Availability of Data and Material

All the data presented in this manuscript are available upon request.

## Author’s Contributions

CP, AB, and MM conceived the experiments. AB and MM collected the data. AB, MM, MB, MK, and CP analyzed and interpreted data, AB and CP wrote the manuscript. All authors reviewed and provided final approval of the version to be published and agree to be accountable for all aspects of the work in ensuring that questions related to the accuracy or integrity of any part of the work are appropriately investigated and resolved. All people designated as authors qualify for authorship, and all those who qualify for authorship are listed. CP is the guarantor for the work and/or conduct of the study, had full access to all the data in the study and takes responsibility for the integrity of data and the accuracy of the data analysis, and controlled the decision to publish.

## Acknowledgements

We acknowledge the technical support of Mr. Jim Sikes and Mr. Bobby Fay from the Arkansas Children’s Nutrition Center Rodent Metabolic Phenotyping Core.

